# QUAS-R: Glutamine (Q) Uptake Assay with Single cell Resolution reveals metabolic heterogeneity with immune populations

**DOI:** 10.1101/2022.09.29.510040

**Authors:** Leonard Pelgrom, Gavin Davis, Simon O’Shoughnessy, Sander Van Kasteren, David Finlay, Linda Sinclair

## Abstract

System level analysis of single cell data is rapidly transforming the field of immunometabolism. However, metabolic profiling of single cells and small populations by flow and mass cytometry is extremely limited by the availability of specific reagents such as antibodies and fluorescently nutrient analogues. Given the competitive demand for nutrients in pathogenic microenvironments including sites of infection, tumours and autoinflammation, there is a need to understand how and when immune cells access these nutrients. Fluorescent-tagging of nutrients is one approach to study nutrient transport but is extremely limited in its usefulness as tagging usually changes the transport characteristics and transporter specificity of the nutrient. Herein, we developed a completely new approach for single cell analysis of nutrient uptake where a fluorophore is attached to a functionalized amino acid after it has been transported across the plasma membrane and is within the cell. This in-cell biorthogonal labelling ensures that *bona fide* transport has been measured. System ASC transporter SLC1A5/ASCT2 transports multiple amino acids, most notably the crucial fuel glutamine, and has essential roles in supporting immune metabolism, signalling and function. This flow cytometry assay allows for rapid, sensitive, and quantitative measurement of SLC1A5-mediated uptake, which we used to interrogate the transport capacity of the complex immune subpopulations within the thymus, at a single cell resolution previously “unreachable”. Taken together, our findings provide an easy procedure to assess which cells support their function via SLC1A5 mediated uptake of amino acids in a sensitive single cell assay. This assay is a significant addition to the single-cell metabolic toolbox required to decode the metabolic landscape of complex immune microenvironments.

## Introduction

Immune cells have a highly dynamic physiology whereby their metabolic and biosynthetic capacity are dramatically regulated in response to immune stimulation^1^. Immune activated T cells, for example, clonally expand and differentiate into effector cells that are capable of producing cytokines and/or cytolytic molecules that help to clear pathogen-infected cells or tumour cells and change once again as the response contracts and resolves to immunologic memory. Other immune cells such as terminally differentiated plasma B cells produce vast quantities of antibodies; up to 10’000 antibodies per second, approximately 800 million antibodies per day. This equates to the equivalent of their own cellular mass per day. These critical functions of diverse immune cells must be fuelled, both in energetic terms as well as provision of raw building materials.

The metabolic processes that support these dynamic changes in immune cells activity and function have become a focus of research since it became clear that metabolic interventions can have real therapeutic benefits^2,3^. Modulating immune cell metabolism has the potential to be used to support beneficial immune function, prevent immune dysfunctional or even restore immune function in pathological settings. For example inhibition of glutamine metabolism within the tumour microenvironment has been shown to benefit antitumour CD8 T cell responses ^4^ One issue that hampers the development of new metabolism-targeting treatments, is that metabolic measurements on whole populations of immune cells cultured in vitro do not reflect immune cell metabolism as it occurs in vivo. Therefore, we need technologies to measure cellular metabolism at single cell resolution either in vivo or directly ex vivo. The lack of such technologies represents a major barrier to the fields understanding of the actual immunometabolic processes occurring in complex immune populations in vivo. Whilst single cell mRNA sequencing can provide valuable information about an immune cell blueprint and identity, it is less accurate in defining cellular metabolism and absolute functions because it is protein expression and protein activity and most importantly substrate availability that dictates these metabolic features.

Protein expression is a highly energy dependent process reliant on the supply of ATP and amino acids and is finely balanced with concurrent cellular protein degradation processes. Cellular uptake of nutrients is essential for the elevated protein synthesis associated with dynamic immune responses; this uptake provides catabolites for energy production and anabolites used for protein synthesis and proliferation. In the context of T cell activation, multiple high-resolution proteomics studies have demonstrated that naïve T cells have low levels of nutrient transporter expression and correspondingly low levels of nutrient uptake^5-7^. A critical early feature of T cell activation is the coordinated increase in the expression of multiple nutrient transporters, including amino acid transporters SLC7A5 and SLC1A5 (also called ASCT2)^8,9^. Indeed, we know that limiting amounts of key nutrients including glucose, methionine, arginine and glutamine alter and dampen the T cell effector responses^6,9-12^.

Therefore, measuring and quantifying nutrient uptake at a single cell resolution directly ex vivo would provide invaluable insight into immunometabolic activity or potential, thereby generating the high-resolution data required to resolve metabolic heterogeneity within complex immune environments. However, to date this has been largely unachievable because conventional nutrient uptake assays involve tracking uptake of radiolabelled substrates into cells, which is not compatible with single cell approaches such as flow cytometry and microscopy. One approach has been the use of fluorophore labelled nutrient analogues as markers for transporter activity. For example, the use of bodipy-labelled lipids, or 7-nitrobenzofurazan-labelled glucose (NBDG) have been used to provide some method for imaging and quantifying nutrient fluxes in cells. However, attaching a fluorophore to a nutrient has the potential to dramatically change the transport characteristics of that nutrient and as a result this approach tends to be largely inaccurate. For instance, fluorophore modified lipids accumulate in different compartments than lipids modified with smaller detectable groups^13^ and a widely used fluorescently tagged glucose (2-Deoxy-2-[(7-nitro-2,1,3-benzoxadiazol-4-yl)amino]-D-glucose, 2NBDG) is not transported by the glucose transporters expressed on immune cells, Slc2a1 and Slc2a3 ^14-16^. For amino acids, various fluorescent analogues have been synthesized. However, the transporter specificity of these have not been ascertained. The only well-characterized reagent for measuring transporter activity has been the naturally fluorescent kynurenine. This tryptophan metabolite provides a convenient method to measure activity of the System L amino acid transporter Slc7a5^17^. However, for most transporters a convenient fluorescent cargo has not been identified.

One approach that has been gaining traction in the study of carbohydrate and lipid studies has been the use of so-called bioorthogonal reagents in combination with cell-compatible click chemistry ^18-20^. Here, a compound, for example a metabolite, is labelled with a functional group that is both small and stable to the conditions found in a cell^21^. Then, at the end of a labelling period, the metabolite is visualised within the cell by the highly selective ligation of a fluorophore that is itself modified with a chemical group that can selectively react with the chemical group that was introduced in the metabolite. This approach has been extensively used to label many cellular components, such as glycans^21,22^, lipids^23,24^, DNA^25^, RNA^26^ in species ranging from bacteria^27^, to cell lines and even whole metazoans, such as zebrafish and mice^28,29^. In 2002, it was reported that amino acids carrying such bioorthogonal groups (in this case azidohomoalanine) could be incorporated into the *E. coli* proteome in place of methionine and be used to ligate a protein with a Staudinger ligation reaction^30^, and later the alkyne-containing homopropargylglycine was used and ligated via a copper-catalysed Huisgen azide-alkyne cycloaddition (CCAAC) reaction^31^. In 2006, Tirrell and co-workers reported the use of this same approach to label the methionines throughout the entire proteome of a mammalian cell line^32^. This approach has since been applied to the study of protein synthesis in many species^33^. Yet, despite the broad application of the BONCAT-approach, the transporter specificity of these two amino acids has never been specified. Whether they are transported by the same route as the unmodified methionine is unknown.

Herein, we investigated whether certain bioorthogonal amino acids can be used to achieve rapid, sensitive uptake measurements for a defined amino acid transporter with single cell resolution. We characterise the uptake of bioorthogonally reactive amino acids azidohomoalanine (AHA) and homopropargylglycine (HPG) and – to our surprise – find it to be through SLC1A5/ASCT2, the major glutamine transporter expressed in immune cells. We then demonstrate that this bioorthogonal approach delivers an accurate and quantitative single cell measure of SLC1A5 mediated amino acid uptake capacity. Furthermore, we demonstrate the ability of this assay to resolve metabolically distinct cells in a complex, multi-population immune scenario through studying the SLC1A5-mediated amino acid uptake capacity of developing T cells in the thymus and of immune populations in the spleen.

## Results

### Activated T cells take up bioorthogonal amino acids

It is well-established that bioorthogonal amino acids can also be taken up by mammalian cells *in vitro*. For instance, azidohomoalanine (AHA) and homopropargylglycine (HPG) have been shown to be taken up by mammalian cells and incorporated into proteins at the positions of methionine^32,34^. HPG contains an alkyne functional group whereas AHA contains an azide functional group (Fig1A). However, the transporters that mediate cellular uptake of these bioorthogonal amino acids are unknown. We first confirmed that immune activated T cells take up bioorthogonal amino acids; HPG and AHA. Unstimulated or immune activated (anti-CD3/CD28) splenic/lymph node derived T cells were incubated with each bioorthogonal amino acid for 15 minutes followed by cell fixation and permeabilization. The abundance of bioorthogonal amino acids inside the cell was detected by ligating azido-AF488 or alkyne-AF488 to the HPG and AHA respectively. Background fluorescence of the azido-AF488 or alkyne-AF488 was determined by using T cells that were not provided with HPG or AHA but were treated equivalently thereafter (Fig.1B). The data show that activated T cells take significantly more HPG or AHA than naïve T cells (Fig1.B,C). There was a notable difference in background staining of the detection azido-AF488/alkyne-AF488 compounds between unstimulated and activated cells. This highlighted the importance of including these conditions in the assay and adjusting the mean fluorescence intensity (MFI) values to evaluate the uptake accordingly: uptake MFI = total MFI - background MFI (Fig 1C). Naïve T cells show very low uptake of either bioorthogonal amino acid (Fig.1B). In contrast, clearly detectable signals for uptake of HPG and AHA were seen in activated T cells when provided with bioorthogonal amino acid concentrations greater than or equal to 10 μM (Fig.1B). Neither of the bioorthogonal amino acids tested were toxic to naïve or activated T cells at any of the concentrations used (Supplementary Fig.1). Next, the uptake was measured over time into activated CD4 and CD8 T cells at either 37°C or 4°C to determine kinetics of the bioorthogonal amino acid transport. Consistent with active transporter mediated uptake, there was a rapid accumulation of biorthogonal amino acids in both activated CD4 and CD8 T cells within 2 minutes that was blocked when the assay was performed a 4°C (Fig.1D,E).

**Figure 1:**
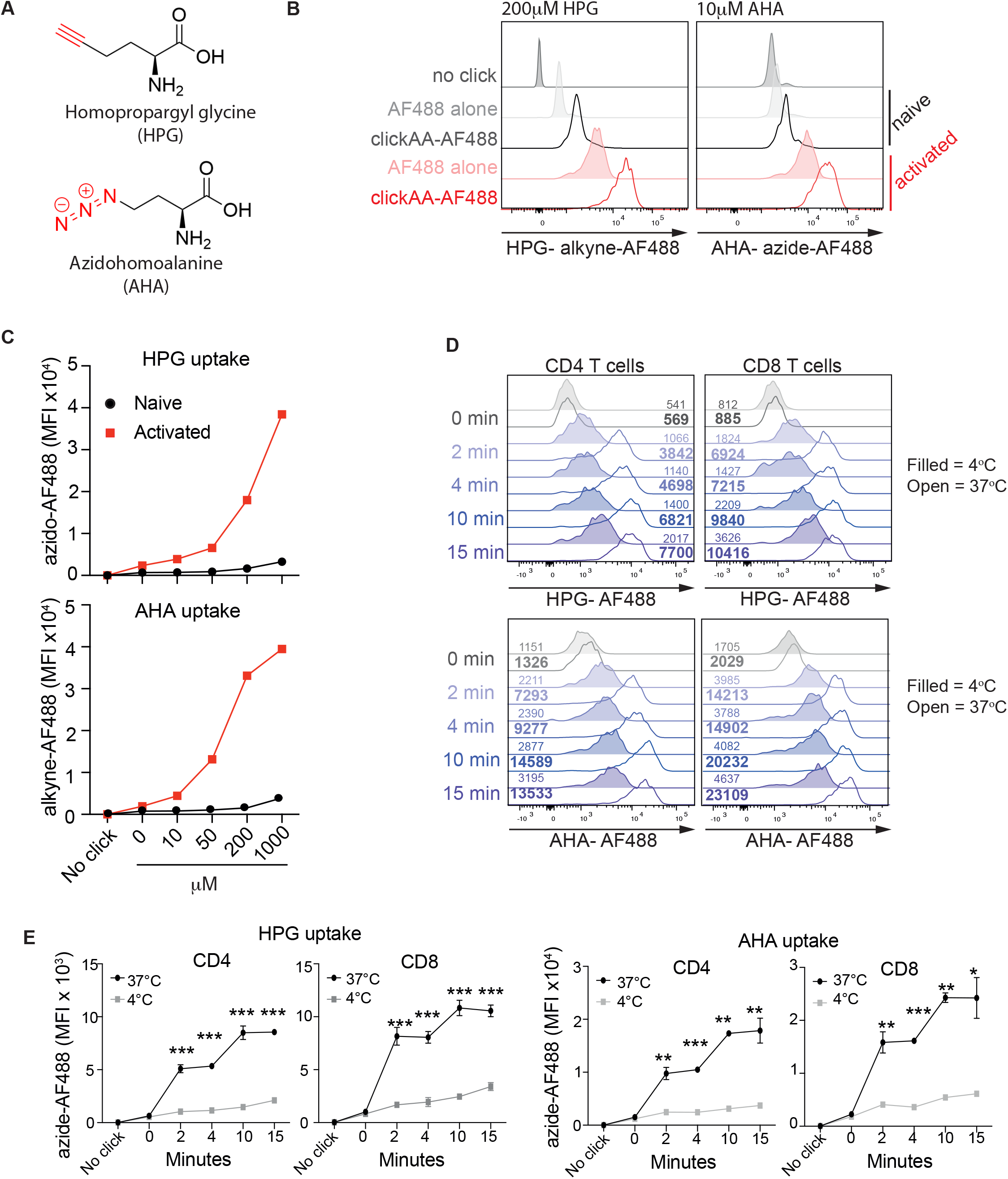
Activated T cells take up bioorthogonal amino acids. **(A)** Schematic representation of the bioorthogonal amino acids Homopropargyl glycine (HPG) and Azidohomoalanine (AHA). HPG contains an alkyne functional group, and AHA contains an azide functional group. The functional groups are indicated in red and are subsequently attached to fluorophores in post-uptake “copper-click” reactions. (**B)** Representative flow cytometry histogram showing HPG (200 μM) or AHA (10 μM) uptake in ex vivo or anti-CD3/CD28 treated (24h) T cells. Uptake performed for 15 mins prior to fixation and subsequent copper-click reaction. No click (HPG or AHA alone), alkyne AF488 alone or azide AF488 alone controls are shown. **(C)** Adjusted uptake MFI (total MFI - background MFI) values for HPG uptake or AHA uptake from ex vivo or anti-CD3/CD28 treated (24h) T cells. Uptake performed for 15 mins with increasing concentrations of HPG or AHA as indicated. **(D)** Representative flow cytometry histogram showing the kinetics and temperature sensitivity of HPG (200 μM) or AHA (10 μM) uptake in anti-CD3/CD28 treated (24h) CD4 and CD8 T cells. Uptakes performed at 4°C indicated by filled histograms, uptakes performed at 37°C indicated by open histograms. Corresponding MFI values are presented on the graph as indicated. **E)** Pooled data showing the adjusted uptake MFI (total MFI - background MFI) values for HPG uptake or AHA uptake as in (D). *P values *** = <* 0.001. Data are representative of at least 3 independent experiments.

In separate experiments the uptake of HPG into type 1 dendritic cells (cDC1), present within murine splenocyte preparations, was measured by both flow cytometry and confocal microscopy (Fig.2AB). Splenocytes were stained with surface antibodies to identify Xcr1+ cDC1 and then provided with HPG for 2 minutes at either 37°C or 4°C, before fixation and permeabilization. Uptake of HPG was visualised as above using an azido-AF488 and the copper-click conjugation reaction. There was a significant HPG uptake signal detected by flow cytometry in cDC1 at 37°C that was prevented when the uptake was performed at 4°C (Fig. 2A). In parallel, uptake visualised and quantified by confocal microscopy and similarly, an uptake signal was measured at 37°C that was inhibited at 4°C (Fig.2B). The uptake of HPG into Xcr1+ cDC1 was found to be significantly greater than that of non-cDC1 splenocytes (Xcr1-cells) both by flow cytometry and confocal microscopy (Fig.2C,D).

**Figure 2:**
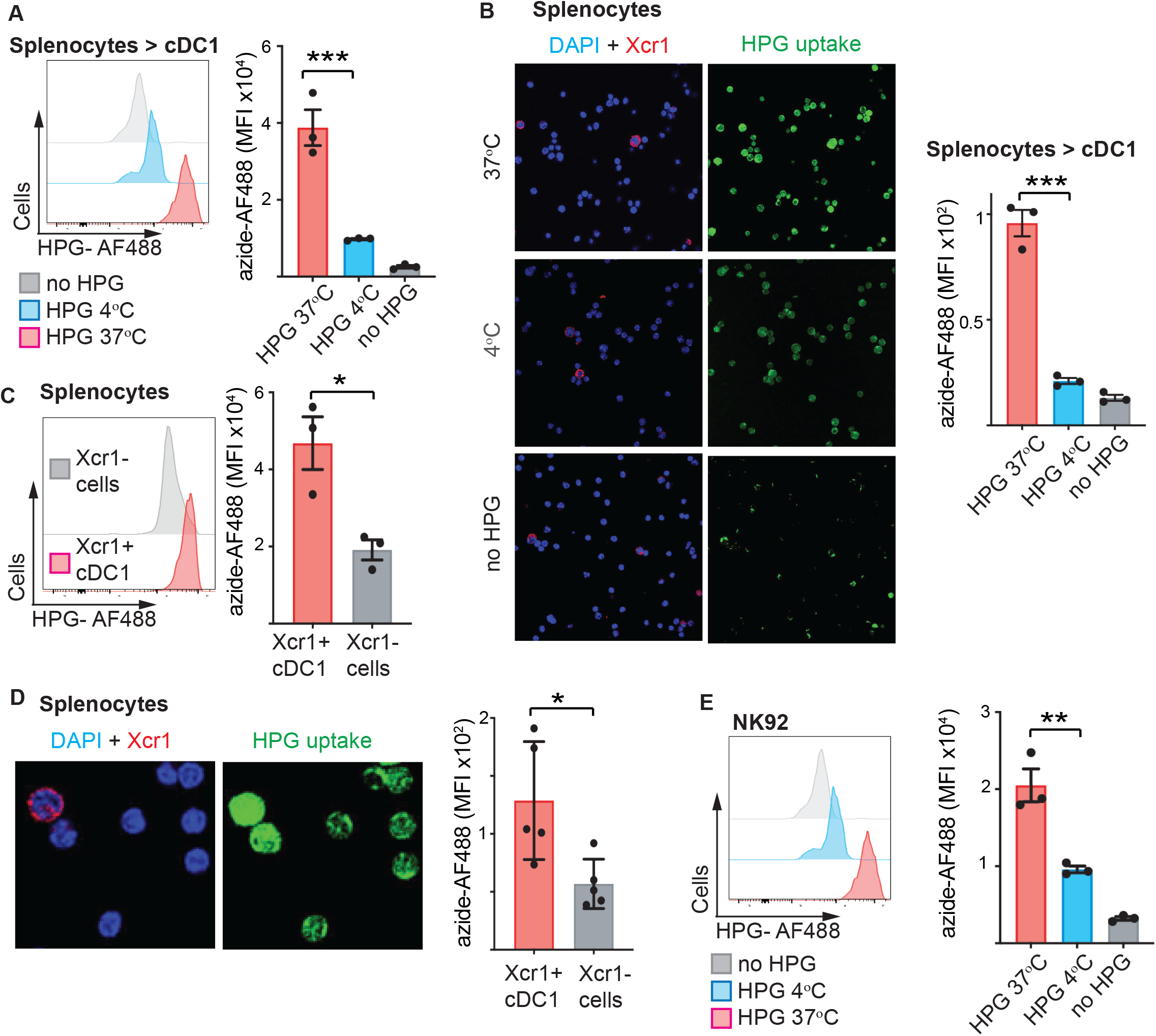
Type 1 Dendritic cells and human NK cells take up bioorthogonal amino acids. **(A)** Representative flow cytometry histogram showing HPG uptake in ex vivo MHCII^high^, CD11c^+^ Xcr1^+^ cDC1 (left) and quantification shown in bar chart on right. Splenocytes were stained with surface antibodies to identify cDC1 prior to 2 min uptake of HPG (200μM) at either 37°C or 4°C. Uptake was stopped with fixation and subsequent copper-click reaction performed. **(B)** Parallel microscopy images of splenocytes as in (A): co-stained with nuclear stain DAPI (blue) and cDC1 marker Xcr1-AF647 (red), with corresponding images showing HPG-AF488 uptake in green. Quantification of pooled data is shown in the bar chart (right). (**C)** Representative flow cytometry histogram (left) and quantification (right) showing HPG uptake at 37°C in Xcr1^-^ and Xcr1^+^ splenocytes. **(D)** Confocal microscopy images of splenocytes stained with nuclear stain DAPI (blue) and cDC1 marker Xcr1 (red), with corresponding HPG uptake in green. Quantification of uptake in Xcr1^+^ cells compared with Xcr1^-^ cells is shown. **(E)** Representative flow cytometry histogram showing HPG uptake in human NK cell line, NK92, (left) and quantification shown in bar chart on right. 2 min uptake of HPG (200 μM) at either 37°C or 4°C was performed, prior to fixation and subsequent copper-click AF488 reaction. *P values * = <0*.*01; ** = <0*.*005; *** = <*0.001. Data are representative of at least 3 independent experiments.

While murine and human amino acid transporters share substantial protein homology, it was nevertheless important to test whether the bioorthogonal amino acid HPG is similarly transported into human immune cells. To investigate this, we used the human NK92 cell line and measured a significant uptake signal when cells were provided HPG for 2 minutes at 37°C and this signal was largely reduced when the uptake was performed at 4°C (Fig.2E).

### HPG and AHA are transported by the glutamine transporter SLC1A5

Activated CD8 T cells and NK cells express multiple amino acid transporters (Fig 3A) and so experiments were designed to define which transporter is responsible for the uptake of HPG and AHA. Amino acid transporters can be identified based upon biochemical/biophysical transport parameters. For example, the System L transporters are sodium-independent whereas substrate transport via System ASC and System N and A transporters (SNAT) are sodium dependent. As previously mentioned, the bioorthogonal amino acids AHA and HPG are incorporated into proteins in place of methionine, which means that they bind to the methionine-specific tRNA for use during protein translation^32,34^. If AHA and HPG are also transported similarly to methionine they would be transported by the sodium-independent amino acid transporter Slc7a5, the primary methionine transporter in activated T cells^6^. It was, therefore, surprising that the uptake of HPG and AHA in to CD8 T cells was sodium dependent (Fig.3B). Uptake was measured in sodium containing and sodium free buffers and the data showed that both HPG and AHA uptake were reduced to the level of the 4°C control in the sodium free buffer (Fig.3,B). Similar data was obtained for activated CD4 T cells (Supplementary Fig.2A). This indicates that transport is not via Slc7a5 but is instead mediated via a sodium dependent amino acid transporter. To further explore what transporter is required for AHA and HPG uptake, competition experiments were performed with amino acids known to be substrates for sodium-dependent transporters. It was found that uptake of both HPG and ALA were inhibited by competition with alanine (Fig 3C). Activated T cells increase the expression of two sodium-dependent alanine preferring transporter families: SLC1 (SLC1A5) and SLC38 (SLC38A1 and A2)(Fig 3A)^35^. Uptake via SLC38A1 and A2 can be distinguished from uptake via SLC1A5 by use of the SLC38 competitive substrate methylaminoisobutyric acid (MeAIB). Hence uptake via SLC1A5 would be unaffected by competition with MeAIB, whereas uptake via SLC38A1 and A2 would be impaired. Figure 3D shows that HPG and AHA uptake is competitively blocked by alanine with an IC_50_-value of 825 μM and 540 μM, respectively. In contrast, MeAIB did not inhibit uptake at any concentration assayed. Lysine was used as a control amino acid that is not a substrate for either SLC1A5 or SLC38A1/2 and also did not inhibit AHA or HPG uptake (Fig 3D,E). Lysine is however a substrate for Y+LAT2/SLC7A6, which is also expressed by T cells albeit at lower levels. Y+LAT2/SLC7A6 is both sodium dependent and independent. It is sodium independent in its capacity to transport arginine, but sodium dependent regarding transport of neutral amino acids including leucine and glutamine^36^. Taken together, these data argue that SLC1A5 is the predominant transporter involved in HPG and AHA transport into activated T cells.

**Figure 3:**
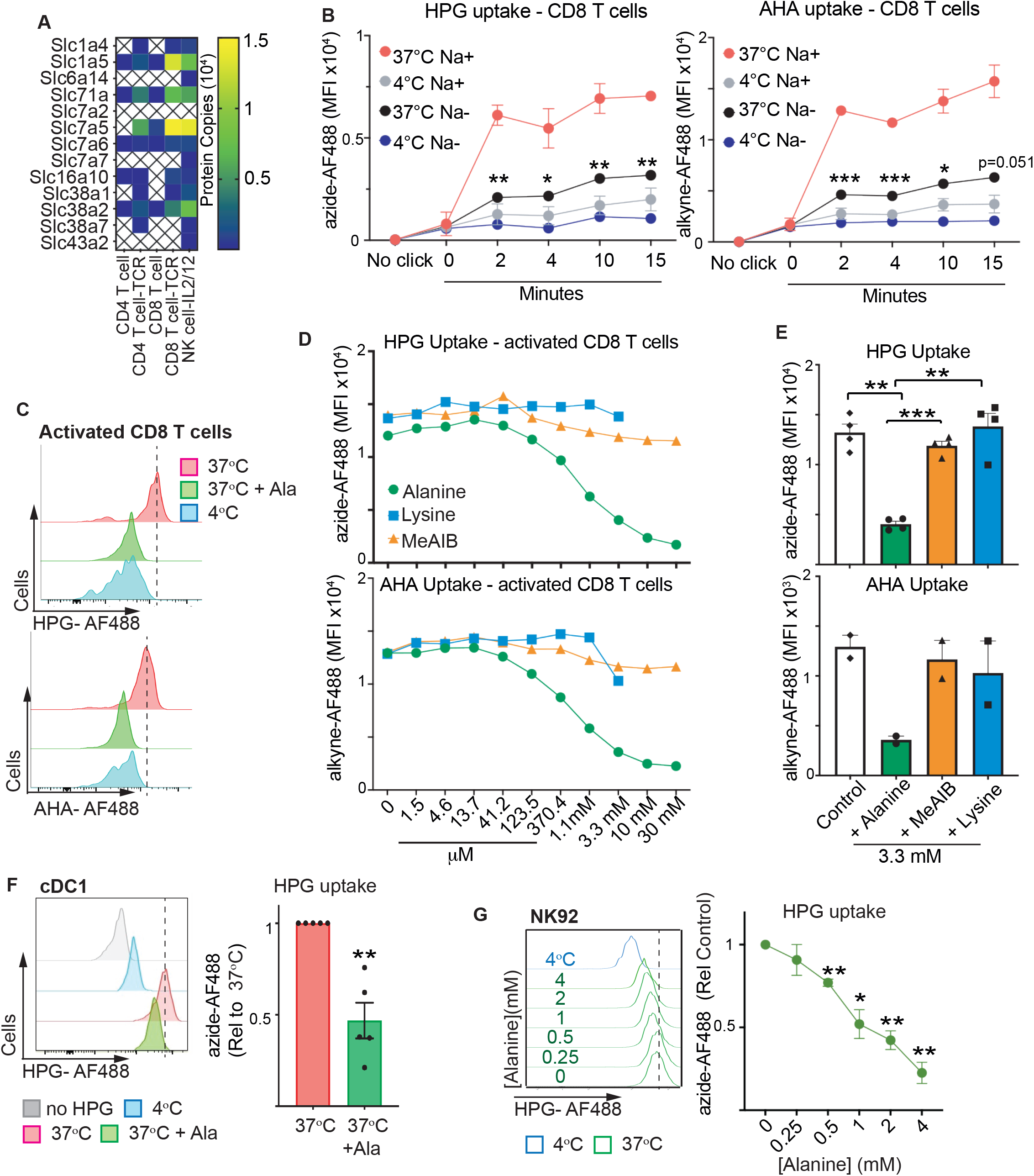
Identification of SLC1A5 as the transporter of HPG and AHA in immune cells. **(A)** Heatmap showing expression of plasma membrane amino acid transporters expressed in murine naïve CD4 T cells, TCR activated CD4 T cells, naïve CD8 T cells, TCR activated CD8 T cells and IL2/12 activated NK cells. Scale bar shows blue (low expression) to yellow (high expression) with corresponding protein copy numbers indicated. Data from Immpres.co.uk. **(B)** Adjusted uptake MFI (total MFI - background MFI) values for HPG or AHA from activated (anti-CD3/CD28 treated; 24h) CD8 T cells in Na+ containing or Na-free uptake media at 37°C or 4°C for indicated times. **(C)** Representative flow cytometry histograms showing HPG (200 μM; 2min; top panel) or AHA (10 μM; 2 min; bottom panel) uptake into activated CD8 T cells in the presence or absence of 5 mM Alanine. Uptake at 4°C control also shown. **(D)** Adjusted uptake MFI (total MFI - background MFI) values for HPG or AHA in activated (anti-CD3/CD28 treated; 24h) CD8 T cells. Uptakes were performed in the presence of increasing concentrations of the competitive substrates Alanine, Lysine or MeAIB. **(E)** Pooled data showing HPG and AHA uptake MFI in the presence of 3.3 mM Alanine, MeAIB or Lysine. Each point indicates an independent experiment. **(F)** Representative flow cytometry histogram (left panel) and quantification bar chart (right panel) showing alanine competition (5 mM) of HPG uptake (200 μM; 2min) into ex vivo spleen derived Xcr1^+^ cDC1. Uptake at 4°C control also shown. **(G)** Representative flow cytometry histograms (left panel) and pooled quantification (right panel) showing HPG (200μM; 2min) uptake into NK92 cells in the presence of increasing concentrations of alanine as indicated. *P values * = <0*.*01; ** = <0*.*005; *** = <*0.001. Data are representative or pooled from at least 3 independent experiments. (D+E) AHA uptake from 2 independent experiments.

To confirm the results obtained in murine T cells, we performed similar uptake experiments for murine cDC1 and human NK92 cells. The data show that competition with alanine reduced the uptake of HPG into cDC1 cells as measured by flow cytometry (Fig.3F). The Immgen RNAseq dataset GSE127267 shows that Slc1a5 is the most highly expressed glutamine transporter in cDC1 (Supplementary Fig.2B). Similarly, RNAseq data from GSE26876 show that SLC1A5 is expressed in NK92 cells (Supplementary Fig.2C). Consistent with SLC1A5-mediated uptake of HPG into NK92 cells, the data showed that the uptake of HPG was inhibited by competition with alanine (Fig.3G).

One important substrate transported by SLC1A5 is the amino acid glutamine, which acts as both a fuel for energy generation and a carbon source for biosynthesis. Indeed, SLC1A5 is the predominant glutamine transporter expressed in many immune cells (Immgen.org and immpres.co.uk)^35^. The data show that AHA and HPG uptake into activated CD8 T cells is also competitively inhibited by increasing glutamine concentrations with an IC_50_-value of <1mM (Fig.4A). This is in line with the Km value of inward glutamine transport for SLC1A5/ASCT2 being in the millimolar range, whereas inward alanine transport is in the μM range^37^. Glutamine also potently competes for HPG uptake into cDC1 cells and NK92 cells providing strong evidence that these bioorthogonal amino acids are accurate reporters of glutamine uptake (Fig.4B,C). The gold standard approach for measuring transporter mediated nutrient uptake is to use radiolabelled substrates. Therefore, we used this approach to provide definitive evidence of SLC1A5-mediated uptake of HPG and AHA. Firstly, we confirmed that both alanine and glutamine would compete with radiolabelled glutamine (^3^H-glutamine) for uptake in activated CD8 T cells. For these radiolabelled uptakes we used in vitro generated IL2 maintained effector CD8 T cells; cytotoxic T cells (CTL). CTL can be expanded into a large, homogenous population which is appropriate for this population-based uptake approach. Furthermore, as with TCR activated T cells, SLC1A5 is the predominant glutamine transporter expressed by IL2 maintained CTL^35^. The data in Figure 4D show that cold glutamine effectively competes for the uptake of ^3^H-glutamine into CTL, with an IC_50_ of 142 μM in this assay, whilst alanine competes ^3^H-glutamine uptake with an IC_50_ of 85 μM. These values are in line with previously published affinities for SLC1A5/ASCT2^37^. Next, we challenged the specificity of HPG and AHA uptake by determining whether each biorthogonal amino acid could act as competitive inhibitor to block the uptake of ^3^H-glutamine. We found that HPG competes ^3^H-glutamine uptake with an IC50 of 386 μM, whilst AHA competes ^3^H-glutamine uptake with an IC50 of 74 μM. In contrast, uptake of ^3^H-glutamine in the presence of the System L competitive substrate 2-aminobicyclo-(2,2,1)-heptane-2-carboxylic acid (BCH) or the SLC38A1/2 substrate MeAIB did not inhibit ^3^H-glutamine uptake (Fig.4E).

**Figure 4:**
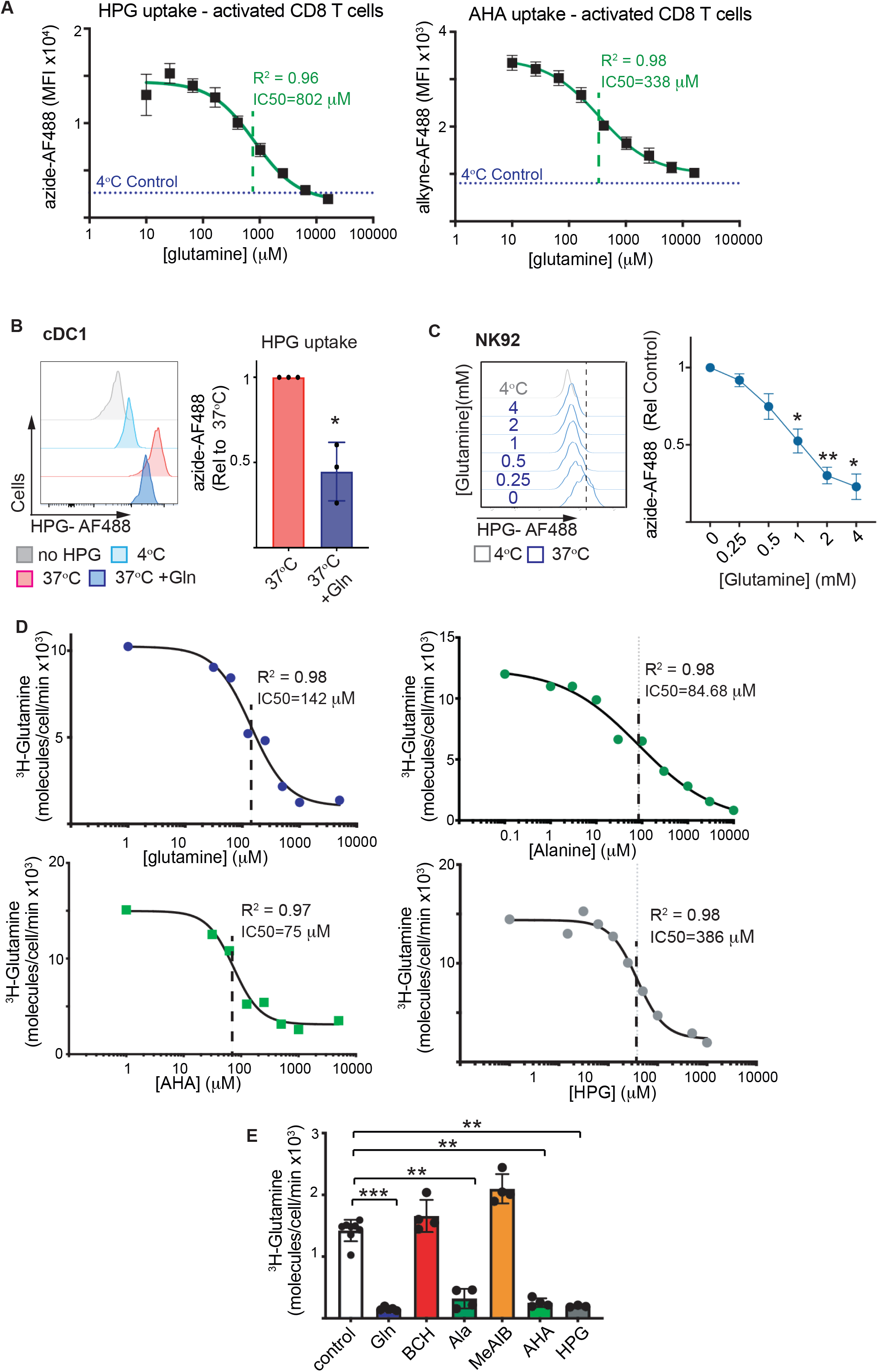
HPG and AHA report Glutamine transport capacity in immune cells. **(A)** Pooled data showing uptake of HPG or AHA in activated (anti-CD3/CD28 treated; 24h) CD8 T cells in the presence of increasing concentrations of Glutamine. Uptake displayed as total MFI - background MFI values. Uptake at 4°C control also shown. **(B)** Representative flow cytometry histogram (left panel) and quantification bar chart (right panel) showing glutamine competition (5 mM) of HPG uptake (200 μM; 2min) into ex vivo spleen derived Xcr1^+^ cDC1. Uptake at 4°C control also shown. **(C)** Representative flow cytometry histograms (left panel) and pooled quantification (right panel) showing HPG (200 μM; 2min) uptake into NK92 cells in the presence of increasing concentrations of glutamine as indicated. Uptake at 4°C control also shown. **(D)** ^3^H-glutamine uptake in effector cytotoxic CD8 T cells (CTLs) in the presence of increasing concentrations of competitive substrates; glutamine, alanine, AHA and HPG. **(E)** Pooled data showing ^3^H-glutamine uptake in effector cytotoxic CD8 T cells (CTLs) in the presence of 5 mM competitive substrates; glutamine, BCH, alanine, MeAIB, AHA and HPG. Data points indicated biological replicates. *P values * = <0*.*01; ** = <0*.*005; *** = <*0.001. Data are representative of at least 3 independent experiments.

Interestingly, we tested another biorthogonal amino acid that we originally predicted would be taken up by SLC1A5 as it was reported to be an isostere of glutamine. This α-azido-glutamine was, however, not taken up by activated T cells. Only at very high concentrations (> 1 mM) could some uptake be seen, which was not competed away with glutamine. Therefore, α-azido-glutamine is neither a reporter for glutamine uptake nor a substrate of SLC1A5 (Supplementary Fig.3A,B).

Taken together, our findings provide an easy procedure to assess which cells support their function via SLC1A5 mediated uptake of amino acids in a sensitive single cell assay, wherein click functionalized reporter molecules, with either an azide or an alkyne group, can be introduced to match the preferred detection method.

### Resolving SLC1A5 uptake capacity in complex immune populations

One barrier to applying this approach for use in complex multiparametric flow cytometry analysis is the fact that copper-click chemistry has the potential to significantly quench the fluorescence of protein fluorophores, such as phycoerythrin (PE), thereby significantly reducing the multiplexing power of this assay^38,39^.

We confirmed that the usual reaction mixture (“click mix”) used for the copper-click reaction did indeed significantly quench PE and PE-Cy7, based fluorescent signals (Fig 5A,B). We found that copper sulfate, especially in combination with sodium ascorbate, led to a loss of PE and PE-Cy7 signal The addition the Cu(I)-stabilising ligand Tris(3-hydroxypropyltriazolylmethyl)amine (THPTA)^40^ and aminoguanidine to the click mix was able to restore fluorescent signals (Fig 5B,C). Aminoguanidine is not required for copper-click, but is sometimes used to prevent unwanted side-reactions^40^. We have previously used it to diminish unwanted side-reactions of the copper in ultrathin cryosections^41^. There were significant increases in fluorescence for PE, PE-Cy7 and PercP-Cy5.5 with increasing concentration of aminoguanidine that plateaued at 160 mM (Fig. 5C). Neither the click mix nor increasing aminoguanidine affect the fluorescent signals for FITC or APC (Fig. 5C). Increasing the concentration of formaldehyde used to fix the cells between antibody labelling and performing the copper-click reaction had no appreciable effect (data not shown), but preservation of the signal worked best when the “click mix” components were prepared fresh for the experiment (Fig. 5D).

**Figure 5:**
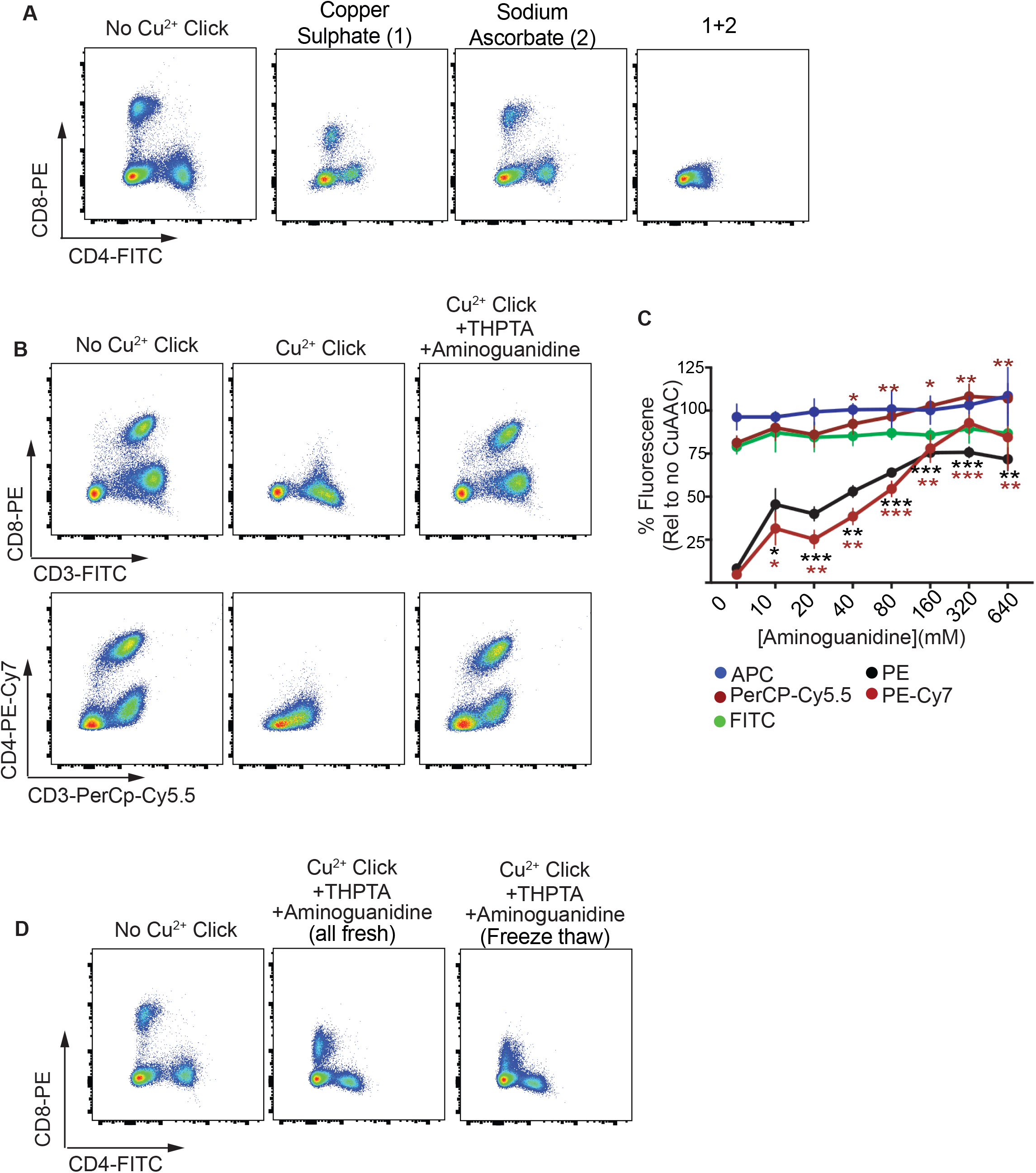
Fine-tuning the Click mix to allow PE/PerCP based fluorophores in antibody panels. **A)** Representative staining of splenocytes with CD4-FITC and CD8-PE, followed by 1 hour treatment with click mix components, copper sulphate (1 mM), sodium ascorbate (10 mM) or both together. **(B)** Representative staining of splenocytes with CD3-FITC and CD8-PE or CD4-PE-Cy7 and CD3-PerCp-Cy5.5 showing the decrease in fluorescence when treated with the click mix for 1 hour and the rescue in fluorescence with the addition of THPTA (1 mM) and aminoguanidine (160 mM). **(C)** Pooled data of fluorescence signals from various fluorophores following incubation with the click mix with THPTA (1 mM) and increasing concentrations of aminoguanidine (1-640 mM). **(D)** Representative staining of splenocytes with CD4-FITC and CD8-PE after incubation with the click mix alone or with THTPA (1 mM) and aminoguanidine (160 mM) that had been prepared fresh or had been freeze-thawed multiple times. *P values * = <0*.*01; ** = <0*.*005; *** = <*0.001. Data are representative (A,B,D) or pooled from at least 3 independent experiments.

Taken together, we believe that the incorporation of aminoguanidine into the “click mix” is an easy and cheap way to improve the compatibility of copper-click with flow cytometry, and now include it in our assay. A simplified overview of the workflow can be found in Supplementary Figure 4.

To challenge the power of our click-based uptake assay to resolve SLC1A5 transporter activity in single cells within a complex immune population, we decided to analyse murine thymocytes directly ex vivo. T cell development occurs in the thymus and involves tightly regulated stages of biosynthesis, proliferation and contraction. These stages are essential to supply a full and varied repertoire of mature T cells which populate the periphery. Conventional thymocyte populations are often categorised according to their expression of the TCR co-receptors CD4 and CD8. Early T cell progenitors entering the thymus from the bone marrow do not express CD4 or CD8 and are termed double negative (DN) thymocytes. During this DN stage, thymocytes that successfully express a functional pre-TCR will undergo robust expansion (DN3-DN4 stage) and differentiate to express both CD4 and CD8 and are termed double positive (DP) cells. A functional mature TCR, alongside either CD4 or CD8 downregulation is required for mature conventional single positive (SP) CD4 or CD8 T cells to exit the thymus and home into the periphery. A mouse thymus consists of approximately 200-300 million cells; 5% are DN, 80% are DP and approximately 15% are SP. The majority of DP thymocytes fail to generate a functional TCR and as result do not receive survival signals and die by neglect. Those that do express a functional TCR, receive survival signals and finally differentiate into single positive CD4 or CD8 expressing T cells which exit the thymus to populate the peripheral immune system.

Very little is known about nutrient uptake of these different thymocyte populations as they develop under these complex conditions. As such we used our bioorthogonal uptake assay to interrogate the SLC1A5-mediated uptake capacity of developing thymocytes. Taking an “overall view”, copper-click uptake of the whole thymus, which would be comparable to uptake data achieved using radiolabelled substrate, there is relatively little HPG uptake signal (Fig.6A, Supplementary Figure 5). However, it is clear from the flow histogram that there is a heterogeneous uptake into total thymocytes; most cells have low HPG uptake, but there is a small proportion that have high HPG uptake which is sensitive to alanine competition and is inhibited at 4°C (Fig.6A, Supplementary Figure 5). Sub-gating into the distinct thymic differentiation stages demonstrates that HPG uptake levels are distinct for the different thymocyte subsets (Fig. 6B,C). Uptake into DN thymocytes is high and heterogeneous, whereas uptake into DP thymocytes is low, consistent with the low metabolic demands of the DP population where the majority of cells are dying (Fig.6B,C, Supplementary Figure 5). SLC1A5-mediated uptake into CD8 SP or CD4 SP was intermediate between DN and DP thymocytes (Fig.6C, Supplementary Figure 5). The heterogeneity within DN thymocytes was further probed, through separating out DN thymocytes based on the expression of CD25 and CD44. Large metabolic changes occur as thymocytes transition from DN3 to DN4 cells where pre-TCR expressing thymocytes undergo robust growth and proliferation^42^. Consistent with the increased anabolic demands that occur at this stage a large increase in SLC1A5 mediated uptake was measured (Fig.6D). Indeed, this analysis resolved clear differences in the SLC1A5-uptake capacity of the developing thymocytes subpopulations that closely aligned to the mRNA expression pattern of Slc1a5 during thymocyte development (Fig.6E, Supplementary Figure 5).

**Figure 6:**
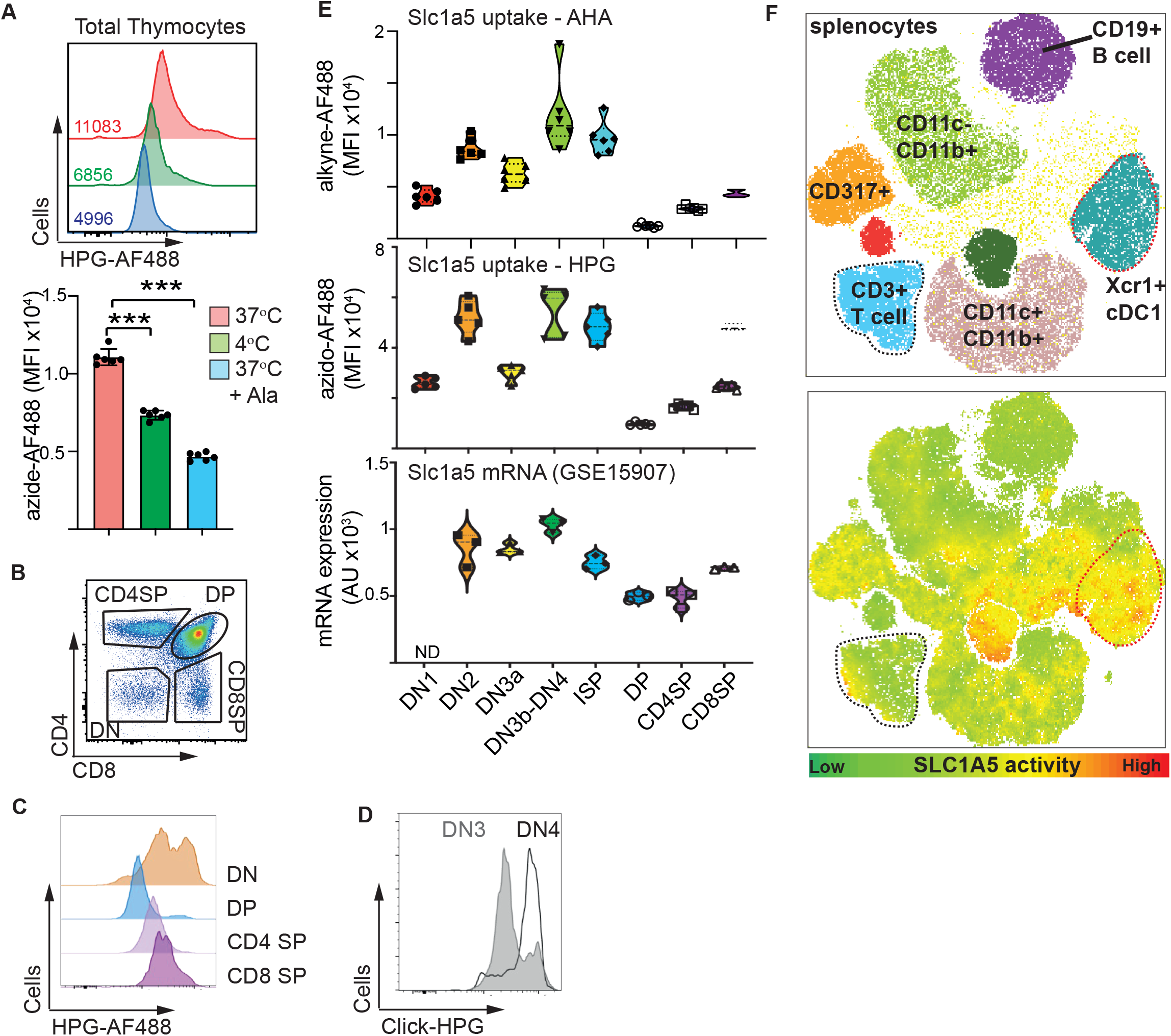
Resolving SLC1A5 uptake capacity in complex immune populations. **(A)** Representative flow cytometry histogram (top panel) and quantitation (bottom panel) showing HPG uptake (200 mM; 2 min; 37°C) in ex vivo thymocytes in the presence or absence of 5 mM Alanine. Uptake at 4°C control also shown. **(B)** Representative flow cytometry dot plot showing CD4 and CD8 staining profiles and gating strategy for thymocyte populations. **(C**,**D)** Representative flow cytometry histogram showing HPG uptake (200 mM; 2 min; 37°C) in DN, DP, CD4SP, CD8 SP (C) and further sub-gated DN3 (CD25^+^ CD44^-^) and DN4 (CD25^-^ CD44^-^) populations (D). (**E)** Pooled data showing HPG (200 mM; 2 min; 37°C; top) and AHA (10 mM; 2 min; 37°C; middle) uptake MFI in thymocytes subsets, aligned with mRNA expression data (bottom panel) from GSE15907 (Immgen.org) for *Slc1a5* from correlating thymocyte subsets. Each data point represents a biological replicate. **(F)** Dimensionality reduction using t-distributed stochastic neighbour embedding (t-SNE) was performed on ex vivo splenocytes treated with HPG (200 μM; 2 min; 37°C). Upper panel shows cDC1 in aqua are gated as CD11c^+^MHCII^+^XCR1^+^CD11b^-^ positive cells; cDC2 in brown are CD11c^+^MHCII^+^XCR1^-^CD11b^+^ cells; pDC in orange are CD11c^int^CD317^+^SiglecH^+^ cells; T cells in blue are CD3^+^CD19^-^ cells; B cells in purple are CD19^+^CD3^-^ cells. Lower panel shows the corresponding HPG uptake for the cell populations. cDC1 are marked with red dotted line, CD3 T cells with a black dotted line. *P values **** = <*0.005. Data are representative of at least 3 independent experiments.

Finally, we probed the glutamine transporter uptake patterns within splenocytes when provided HPG directly ex vivo and combined with multiparametric flow cytometry to identify immune cell populations. The staining panel used identified T cells^CD3+^, B cells^CD19+^ and multiple antigen presenting cells including CD317+ pDC, monocytes^CD11c-/CD11b+^, DC^CD11c+/CD11b+^ and cDC1^Xcr1+^

The data show distinct patterns of HPG uptake with the highest levels identified in the cDC1^Xcr1+^, as well as several unknown populations – which were not identified by the antibody panel used (Fig.6F). The data in figure 6 highlight the potential for using bioorthogonal amino acid uptake assays to unveil metabolic features of ex vivo immune cells that were, until now, unreachable.

## Discussion

It is now clear that the metabolic features of immune cells are substantially altered after in vitro culture compared to those of immune cells in vivo^43^. This highlights the importance of studying immune metabolism in vivo or directly ex vivo. However, the lack of robust technologies to measure metabolic fluxes at a single cell level represents a barrier preventing such in vivo, or directly ex vivo analysis. This barrier is limiting our understanding of the metabolic demands, constraints, and heterogeneity within complex immune populations.

The research described herein provides a robust assay for the amino acid transport through SLC1A5, a transporter known to be important for the metabolism of numerous immune subsets. Through using multiple assays including radiolabelled uptake competition, Na+ dependence as well as substrate competition, we show that the bioorthogonal amino acids AHA and HPG are taken up by primary murine T cells through SLC1A5. We also confirm that AHA and HPG are transported in the human NK cell line, NK92 cells, and in murine type 1 dendritic cells. This assay uses a novel approach wherein the fluorophore is attached to the bioorthogonal amino acids after it has been transported into the cell using “click” chemistry. This innovative approach underpins the accuracy of this assay and avoids the pitfalls common to other nutrient uptake assays, such as 2NBDG. It should be noted that some immune cells, such as immature cDC1 cells, actively uptake material through transporter independent mechanisms such as pinocytosis. Indeed, it is worth noting that the level of HPG uptake in the presence of competition with alanine or glutamine does not come down to that of the 4°C control. In this case the difference in fluorescence between the HPG uptake +/- the competition controls correspond to SLC1A5 uptake. Whereas the difference in fluorescence between the alanine or glutamine competition control and the 4°C control most likely reflects uptake by transporter-independent mechanisms including pinocytosis. This highlights the importance of including the correct controls for all experiments.

Harnessing the single cell resolution of this metabolic assay has allowed, for the first time, the measurement of SLC1A5 amino acid flux in the various thymocyte subsets developing in the thymus. These metabolic fluxes were found to correlate closely with the expression pattern of Slc1a5 mRNA shown in the Immgen dataset GSE15907. Clear heterogeneity in glutamine uptake capacity was also measured within the immune subsets within the spleen directly ex vivo. This showed that type 1 dendritic cells were amongst the cells with the highest uptake whereas most other splenic lymphocyte populations showed low levels of HPG transport. Interestingly, even within immune subsets in these naive mice there was heterogeneity including a small but distinct subset of T cells with elevated HPG, and hence glutamine transport capacity. Together this data demonstrates the power of this novel metabolic analysis in identifying differences in the metabolic activities of immune cells in complex populations.

During the course of this investigation, we also refined the click mix required for the conjugation of the detection fluorophore to the transported bioorthogonal amino acid. This now allows for full multiplexing including the use of PE based fluorophores, previously difficult to achieve.

This work has validated two distinct bioorthogonal amino acids as substrates for SLC1A5. AHA contains an azide to allow it to be visualised using an alkyne-fluorophore while HPG contains an alkyne and can be visualised using an azide-fluorophore. This provides flexibility where SLC1A5 flux analysis can be multiplexed to other assays that utilise bioorthogonal chemical reactions with a different bioorthogonal structure. This would involve a double copper-click reaction using two separate fluorophores, one with an azide and the other with an alkyne. For instance, *azide* containing AHA uptake can be combined with Edu incorporation (measuring DNA synthesis with an *alkyne*-nucleotide analogue) or puromycin incorporation (measuring protein translation using an *alkyne*-puromycin molecule). Similarly, as additional assays of nutrient transporters become available, using this bioorthogonal strategy, it will become possible to measure two nutrient uptake fluxes simultaneously into a single cell. Altogether, our Glutamine (Q) Uptake Assay with Single cell Resolution (QUAS-R) represents a powerful addition to the ever-growing “single-cell metabolic toolkit”.

## Figure Legends

**Supplementary Figure 1:**
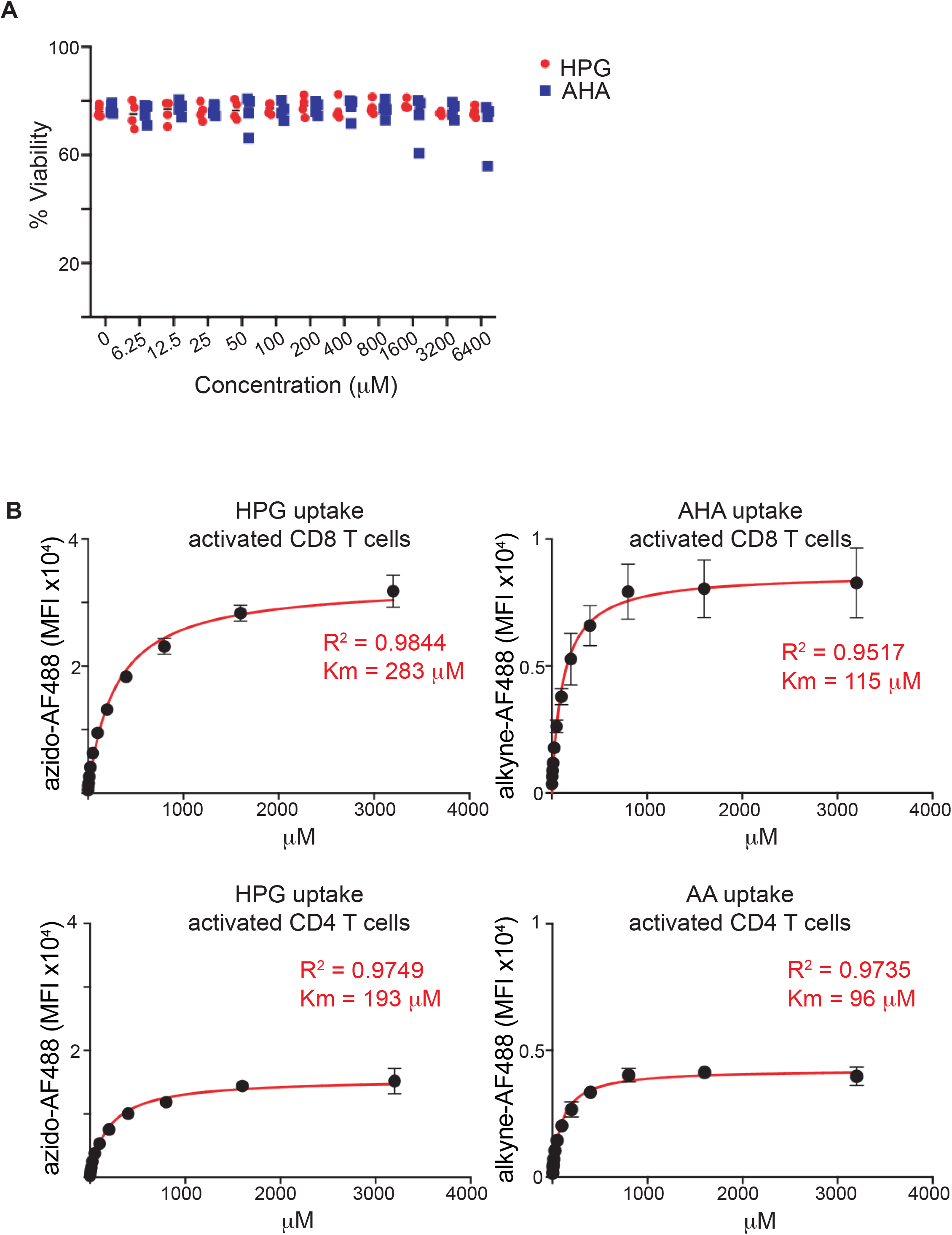
**(A)** Data shows the viability of activated T cells that had been cultured with HPG or AHA in HBSS at the concentrations shown for 15 min, fixed and analysed by flow cytometry using a viability dye. **(B)** Activated CD8 (top) or CD4 (bottom) were cultured with HPG (left) or AHA (right) at increased concentrations for 2 min, then fixed, permeabilised and azido-AF488 (left) or alkyne-AF488 (right) attached via a copper-click reaction. Cells were analysed by flow cytometry and mean fluorescence intensity (MFI) plotted against the concentration of HPG or AHA. The data was fit to a hyperbolic curve and R^2^ and Km values calculated from the resultant graph. Data is from 4 separate experiments (A) and 1 experiment performed in quintuplicate (B).

**Supplementary Figure 2:**
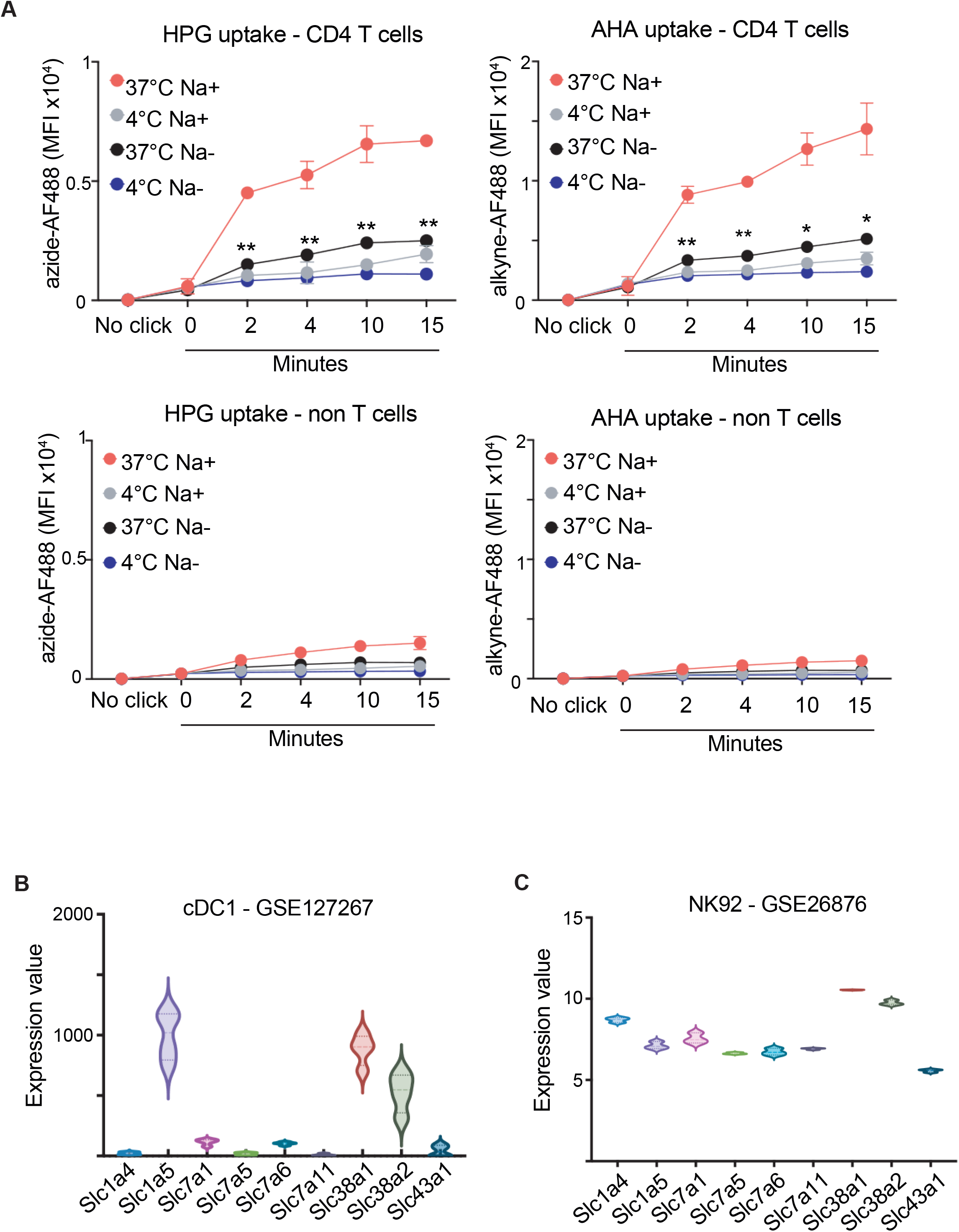
**(A)** Activated T cells cultures were incubated with HPG (200 μM, left) or AHA (10 μM, right) for 0 to 15 minutes in sodium containing and sodium free buffer at either 4°C or 37°C. Cells were then fixed, permeabilised and azido-AF488 (left) or alkyne-AF488 (right) attached via a copper-click reaction. Cells were analysed by flow cytometry and mean fluorescence intensity (MFI) for CD4 T cells (top) and non-T cells (bottom) graphed. **(B**,**C)** RNAseq data sets (GSE127267 and GSE26876) were investigated for the relative expression of mRNA for glutamine amino acid transporters in cDC1 (B) and NK92 cells (C). *P values * = <0*.*05; ** = <0*.*01*. Data are pooled from at least 3 independent experiments.

**Supplementary Figure 3:**
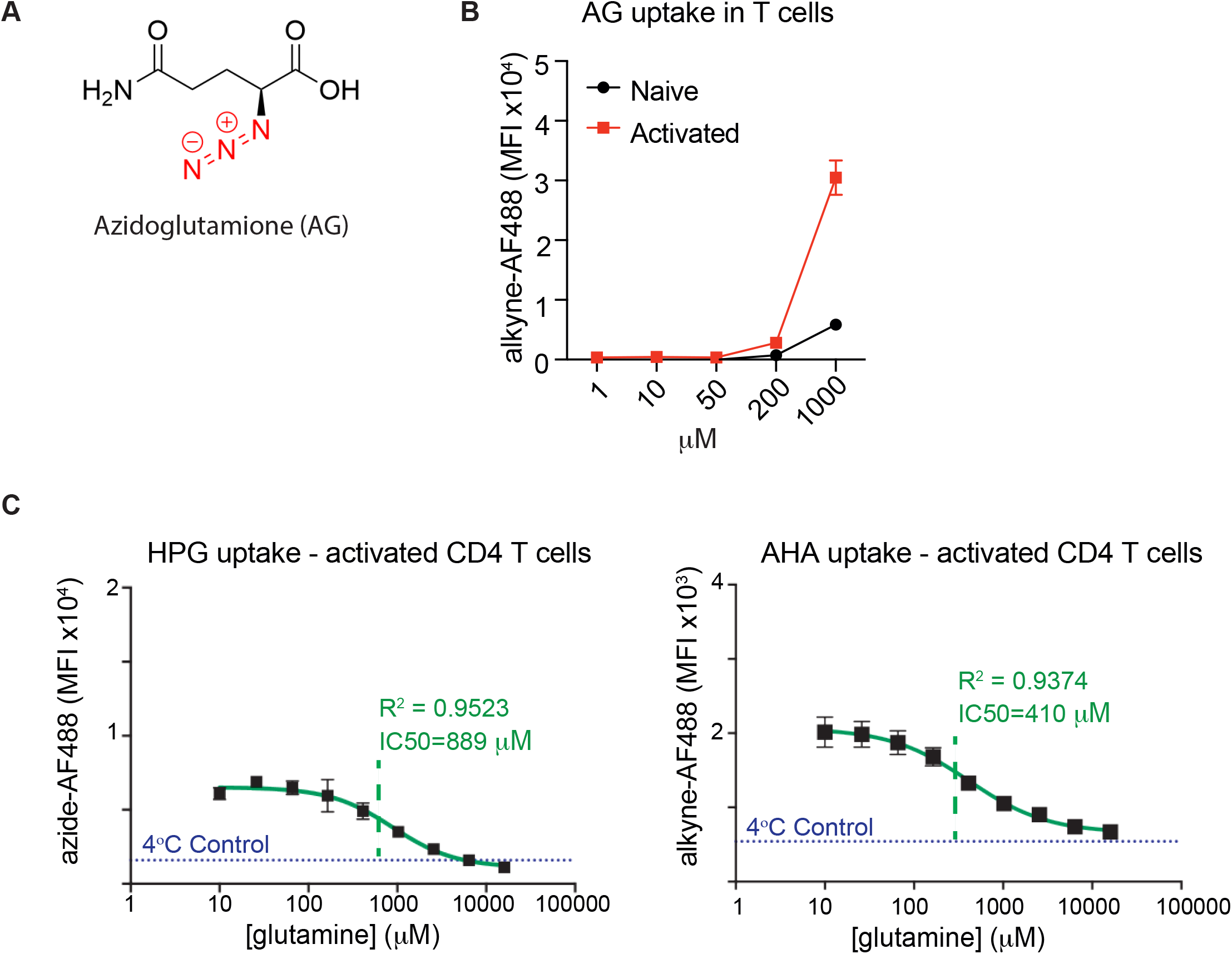
**(A)** Structure of azido-glutamine (AG) is shown. **(B)** Naïve or activated T cells were incubated with AG and increasing concentrations for 15 min at 37°C. Cells were fixed, permeabilised and alkyne-AF488 attached via a copper-click reaction. Cells were analysed by flow cytometry and mean fluorescence intensity (MFI) for T cells was graphed. **(C)** T cells were activated (anti-CD3/CD28 treated; 24h) and then incubated with HPG (200 μM, left) or AHA (10 μM, right) for 2 min at 37°C in the presence of increasing concentrations of glutamine. Uptake of 4°C control also shown. Cells were then fixed, permeabilised and azido-AF488 (left) or alkyne-AF488 (right) attached via a copper-click reaction. Cells were analysed by flow cytometry and mean fluorescence intensity (MFI) in CD4 T cells was plotted against the concentration of glutamine added. The data was fit to a sigmoidal curve and R^2^ and IC50 values calculated. Data is representative of at least 3 independent experiments.

**Supplementary Figure 4:**
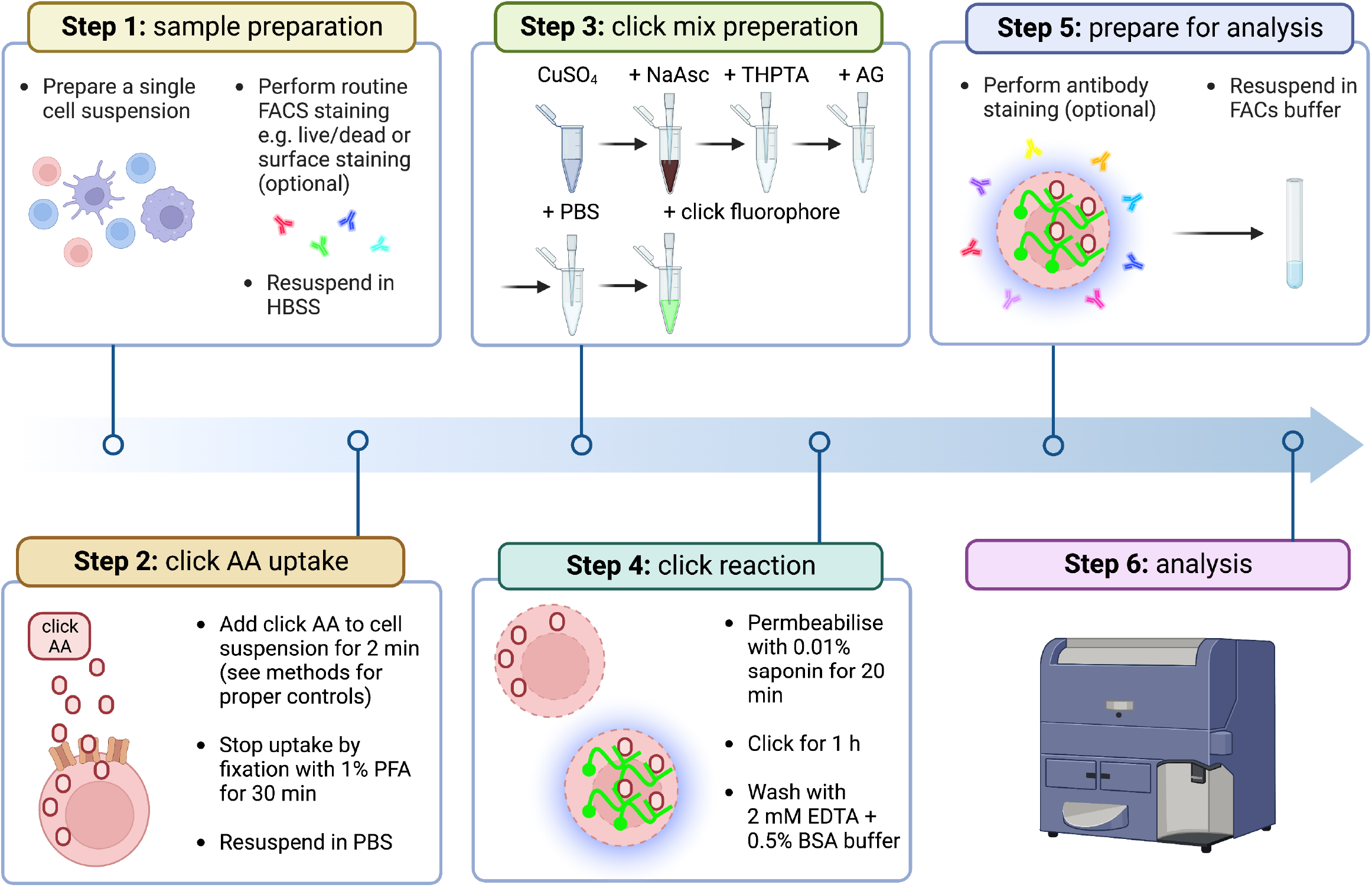
Diagram outlining the key steps for QUAS-R.

**Supplementary Figure 5:**
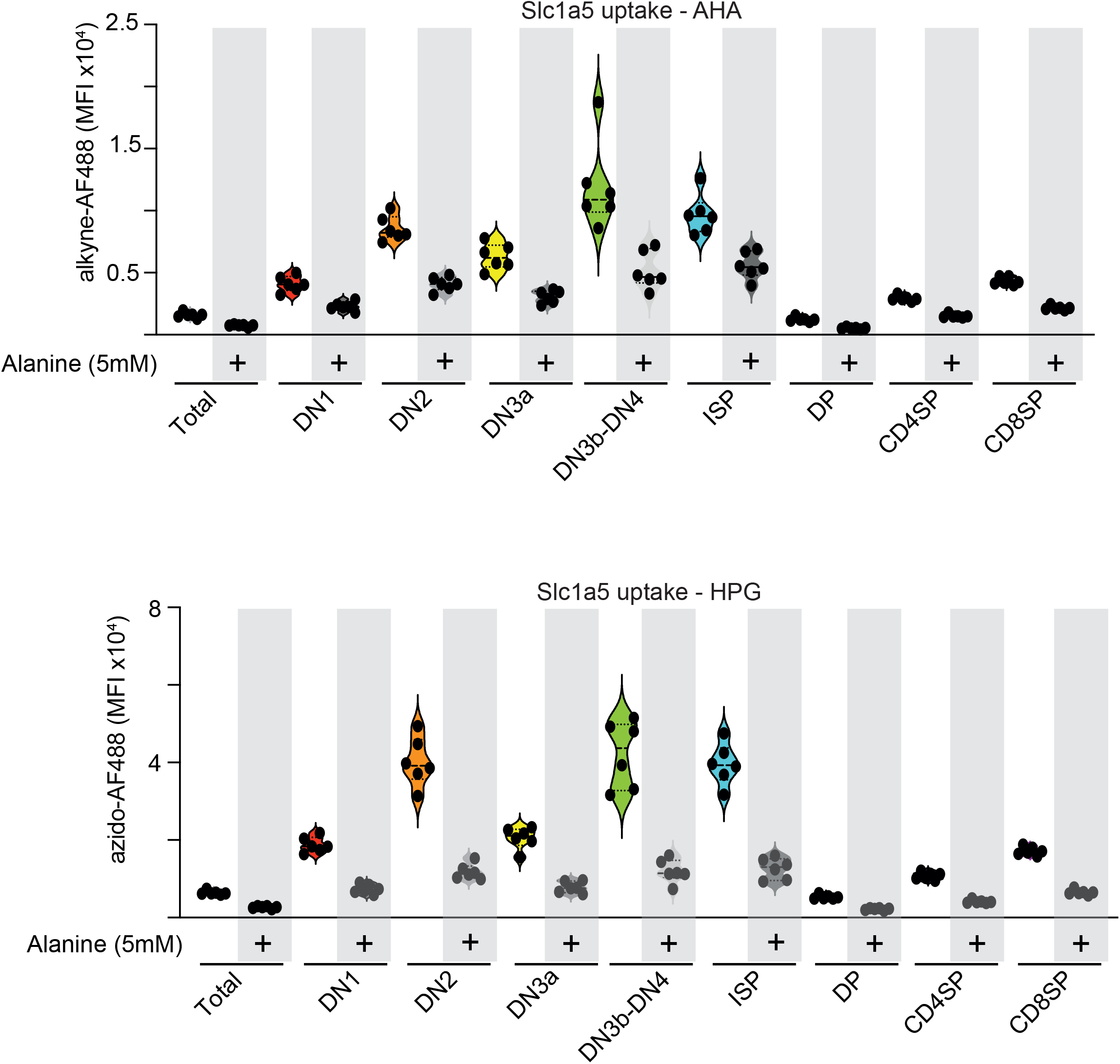
Thymocytes were isolated and cultured with AHA (10 μM, top) or HPG (200μM, bottom) for 2min at 37°C in the presence or absence of alanine (5 mM). Cells were then fixed, permeabilised and alkyne-AF488 (top) or azido-AF488 (bottom) attached via copper click reaction. Cells were analysed by flow cytometry and mean fluorescence intensity (MFI) for each thymocyte population was plotted. Data shown as violin plots with individual data points and pooled from at least 3 independent experiments.

## Materials and methods

### Mice

At the Leiden Institute of Chemistry (LIC), C57BL/6J male and female mice were bred under specific pathogen free (SPF) conditions. Animal license number AVD1160020198832. Mice between 6 weeks and 6 months old were culled by cervical dislocation. Animal experiments were approved by the Dutch Central Authority for Scientific Procedures on Animals (CCD) and performed in accordance with European Union Directive 2010/63EU, Recommendation 2007/526/EC and local government regulations.

At the University of Dundee, OT-1 transgenic mice, whose T cell receptor (TCR) was designed to recognize OVA_257-264_ peptide (SIINFEKL), were bred and maintained in the Biological Resource Unit under SPF conditions. Procedures were approved by the University Ethical Review Committee and under the authorisation of the UK Home Office Animals (Scientific Procedures) Act 1986. Male and Female mice used were between 8 and 14 weeks old.

In Trinity College Dublin, C57BL/6J mice were bred in house and used between 6 and 12 weeks of age. Mice were housed under 12:12 light cycle in a relative humidity of 45-65% and a temperature between 20-24°C. Mouse experiments were approved by and in compliance with the Irish Health Products Regulatory Authority (Project licence AE19136_P091) and the Animal Research Ethics Committee (AREC) at Trinity College Dublin.

### T cell isolation and activation

Spleen from C57BL/6J mice were mechanically disrupted using the back-end of a syringe before addition of 50 μL 11x concentrated collagenase D (Roche, #11088866001; end concentration = 1 mg/mL) and DNase I (Sigma, #D4263; end concentration = 2000 U/mL) for 20 mins at 37°C, 5% CO_2_. After collagenase digestion, the cell suspension was filtered through 70-100 μm sterile filters and centrifuged. Red blood cells were lysed from the splenocyte suspension using red blood cell lysis buffer (Gibco, #A1049201) for 5 minutes at room temperature. The resultant splenocyte cell suspension was resuspended in RPMI (Sigma, #R5886) supplemented with GlutaMAX (Gibco, #35050061), 10% heat-inactivated fetal calf serum, 50 μM of β-mercaptoethanol (Gibco, #31350), 50 IU/mL penicillin and 50 μg/mL streptomycin: (complete RPMI). To activate primary T cells, single-cell spleen suspensions from C57BL/6J mice were cultured at 5 million cells/ mL in complete RPMI with 1 μg/mL anti-CD3 (BioLegend, #100340) and 1 μg/mL anti-CD28 (BioLegend, #102116) for 24h. Naive T cells were seeded from the same spleen suspensions and kept under the same conditions but without anti-CD3/28.

For generation of cytotoxic T cells (CTLs), single-cell suspensions were generated by mashing lymph nodes and/or spleens from OT-1 mice through a 70 μm strainer. Red blood cells in splenocyte suspension were lysed with 150 mM NH_4_Cl, 10 mM KHCO_3_ and 110 μM Na_2_EDTA in mQ water (pH = 7.8). Single-cell suspensions were activated using 10 ng/mL OVA_257-264_ peptide (SIINFEKL peptide; in house) at a density of 5 million cells/mL and cultured at 37°C with 5% CO_2_ in RPMI 1640 containing glutamine (Invitrogen), supplemented with 10% FBS (Gibco), penicillin/streptomycin (Gibco) and 50 μM β-mercaptoethanol (Sigma) with cytokines IL-12 (2 ng/ml; Peprotech) and IL-2 (20 ng/ml; Proleukin Novartis). Cells were activated for 36 h, washed out of activation media and then expanded for a further 3-4 days in media supplemented with 20 ng/ml IL-2.

### In vivo DC expansion

Flt3l-producing B16 melanoma cells were used to expand the DC population in vivo as previously described^44^. Briefly, 2.5 million melanoma cells in 100 μL PBS were injected subcutaneously into the right flank of the mouse. After 10 days of tumor growth, the mouse was culled and the spleens processed as described above.

### NK92 cell culture

NK92 (ATCC CRL-2408) cells were cultured in RPMI 1640 supplemented with 2 mM glutamine (Gibco), 12.5% horse serum (Sigma), 12.5% foetal calf serum (Biosera), 0.02 mM folic acid (Sigma), 0.2 mM myoinositol (Sigma), 1% penicillin-streptomycin (Gibco) and 0.1 mM β-mercaptoethanol (Gibco). Cells were sub-cultured 1:2 every 2 days and fresh growth medium applied. NK92 cells were gently triturated with a pasteur pipette, counted and seeded in 96-well U-bottomed plates at 0.5 million cells/well before further experimentation.

### Bioorthogonal amino acid uptake by flow cytometry

Maintain the temperature of the cells and buffers between 37°C and room temperature, except for those used for uptake at 4°C (cold controls).

Cells were washed using equal parts HBSS (Gibco, #24020) and RPMI (HBSS:RPMI+GlutaMAX), and centrifuged again. After removal of supernatant, the cell pellet was resuspend in HBSS:RPMI+GlutaMAX and counted. Cells were pre-stained with LIVE/DEAD™ Fixable Aqua Dead Cell Stain (Thermo, #L34957) in the dark for 20 minutes at room temperature. 500’000 cells in HBSS were seeded into v-bottom 96-well plate wells.

In parallel, 80 μL of a HBSS solution with twice the concentration of bioorthogonal amino acid as indicated in the graphs (+/- control competing amino acids as indicated) were added to 96-well plate wells and either warmed in a cell incubator (37°C) or chilled on ice (4°C = cold controls) for 15 minutes to equilibrate the temperature. 50 μL of this click AA 2x concentrate was then added to 50 μL of cells and plates were returned to the incubator or ice to incubate for the designated amount of time. Afterwards, 100 μL of 2% paraformaldehyde solution (Polysciences, #18814-20; methanol free; final concentration = 1%) in HBSS was added to each well and the plates were incubated in the dark for 30 minutes at room temperature. Plates were centrifuged, supernatant was removed, and cell pellets were resuspended in phosphate-buffered saline (PBS). Cells were kept in this PBS and at 4°C for later click chemistry and flow cytometry antibody staining.

For determining sodium dependence of amino acid uptake, buffers were prepared with 2 mM KCl (Sigma, # 60128), 1 mM CaCl_2_ (Merck, #1.02389), 1 mM MgCl_2_ (Ambion, #AM9530) and 10 mM (HEPES; inhouse; pH = 7.5) in MQ water and with either 100 mM NaCl (TJ Baker, #277) for the sodium-containing buffer or 100 mM tetramethylammonium chloride (TMACl) for the sodium-free buffer.

Azidohomoalanine (AHA) and homopropargylglycine (HPG) were synthesized as described previously^45,46^. Azidoglutamine was bought From Chiralix (#CX57717).

### Click chemistry

Cells were permeabilized for 20 minutes at room temperature using 0.01% saponin (Sigma, #47036) in PBS after the plate was centrifuged and supernatant was removed. Near the end of permeabilization the ‘click mixture’ was prepared.

In a sequential order sodium ascorbate (NaAsc), tris-hydroxypropyltriazolylmethylamine (THPTA), aminoguanidine, PBS and bioorthogonal fluorophore were added to copper sulphate (CuSO_4_). This together is called the ‘click mixture’ and is used for incubating the cells. Upon combination of light blue CuSO_4_ and the light yellow NaAsc, the solution turns dark brown/black. Upon subsequent addition of the THPTA, the solution turns pale yellow/colourless. We used 1 part of CuSO_4_ (Sigma, #209198; stock 100 mM in Milli-Q [MQ] water, final concentration 1 mM) together with 1 part of NaAsc (Sigma, #A7631; stock 1 M in MQ, final concentration 10 mM), 1 part of THPTA (stock 100 mM in MQ, final concentration 1 mM), 1 part of aminoguanidine (Cayman Chemicals, #81530; stock 1 M in MQ, final concentration 10 mM), 96 parts of PBS and 0.25 part of bioorthogonal fluorophore (AZDye 488 alkyne, AZDye 488 azide, AZDye 647 alkyne, AZDye 647 azide; all Click Chemistry Tools, #1277, #1275, #1301 and #1299 respectively; 2 mM in DMSO, final concentration 5 μM).

After permeabilization, the plate was centrifuged, supernatant was removed, and cell pellets were resuspended in 30 μL of click mixture and incubated in the dark for 1 hour at room temperature. The plate was centrifuged, copper-containing click mixture supernatant was discarded in accordance with the institute recommendations for aqueous copper disposal, cell pellets were resuspended in 200 μL of PBS supplemented with 0.5% bovine serum albumin [BSA; Sigma, #A9467] and 2 mM ethylenediaminetetraacetic acid [EDTA; from inhouse stock of 0.5 M at pH = 8.0] (FACs buffer) and incubated for 30 minutes in the dark. The plate was centrifuged, any residual copper-containing supernatant was discarded in the same manner as before and cells were washed one more time with FACS buffer before antibody staining and normal washes and supernatant removal and acquisition on the flow cytometer. THPTA was synthesized as described earlier ^40^.

### Flow cytometry

Antibody staining was done in the dark for 30 minutes at 4°C in 30 μL of FACS buffer with antibodies and anti-CD16/32 (TruStain FcX™ PLUS; BioLegend, #156604; 1:100). Acquisition was done on a BD FACSCanto™ II or BD LSRFortessa (both BD Biosciences). Analysis, including dimensionality reduction by t-distributed stochastic neighbor embedding (tSNE), was done using FlowJo (TreeStar, version 10).

### Flow cytometry antibodies

#### Spleen T cell panel

B220 - PerCP-Cyanine5.5 (BioLegend, #103236; 1:400), CD4 – PE-Cyanine7 (BioLegend, #100422; 1:400), I-A/I-E (MHC class II) – APC (BioLegend, #107613; 1:4000) and CD8a – APC-Cyanine7 (BioLegend, #100714; 1:400) was combined with AZDye 488 alkyne/azide. When AZDye 647 alkyne/azide was used, MHCII – APC was replaced by MHCII – FITC (eBioscience, #11-5321-82; 1:2000). In this panel, LIVE/DEAD™ Fixable Aqua was stained before the other markers using a 1:400 dilution.

#### Spleen cDC panel

CD3 – APC (Biolegend, #100312; 1:200), CD19 – PE-Cyanine7 (Biolegend, #552854; 1:200), F4/80 – AF700 (Biorad, #497A700; 1:200), I-A/I-E (MHCII) (EBiosciences, #47532182; 1:400), CD11c – Brilliant Violet 421 (BD, #101257; 1:200), CD8a - PerCP-Cyanine5.5 (BD, #551162; 1:200), CD11b – Brilliant Violet 605 (Biolegend, #101257; 1:200), XCR1 – Brilliant Violet 785 (Biolegend, #148225; 1:200) and CD317 – Brilliant Violet 650 (Biolegend, #127019; 1:200) combined with AZDye 488 alkyne/azide.

#### Thymocyte panel

CD24 – Brilliant Violet 421 (BioLegend, #101825; 1:400), Lineage cocktail – PE (Nk-1.1 – PE [BD Biosciences, #557391; 1:100], Gr-1 – PE [eBioscience, #12-5931-82; 1:400], CD19 – PE [eBioscience, #12-0191-83; 1:100], CD11b – PE [eBioscience, #12-0112-81; 1:800]), CD25 – PerCP-Cyanine5.5 (BD Biosciencs, #561112; 1:100), CD44 – PE-Cyanine7 (eBiosciences, # 25-0441-81; 1:600), CD4 – APC (eBiosciences, #17-0042-83; 1:200) and CD8a – APC-Cyanine7 (BioLegend, #100714; 1:400) combined with AZDye 488 alkyne/azide. In this panel, LIVE/DEAD™ Fixable Aqua was stained before the other markers using a 1:400 dilution.

#### Confocal microscopy

Confocal microscopy was performed using a Leica SP8 scanning confocal microscope. After 10 minutes of FC block (BD Biosciences, 1:100), splenocytes were stained for 30 minutes at room temperature with anti-mouse XCR1 conjugated to AF647 (BioLegend) to identify XCR1^+^ cells. Splenocytes were then washed with PBS, resuspended in HBSS for uptake assays and allowed to reach temperature either in a 37°C incubator or on ice. The uptake assay was performed as described above. After HPG uptake, cells were immediately fixed with 1% PFA for 30 minutes at room temperature. Splenocytes were permeabilised with PBS containing 1% BSA and 0.01% saponin. The click reaction was performed as described above using azide-AF488 in the click mixture. Cells were washed 4 times with PBS and resuspended in PBS containing 1% BSA and DAPI (1 μg/mL; Sigma) as a nuclear stain. Splenocytes were stained for 45 minutes with DAPI before washing. Finally, cells were resuspended in a mounting medium (Mowiol, Sigma) and pipetted directly onto high-resolution microscope slides. The mounting medium was allowed to set overnight at room temperature in the dark before imaging. Quantification of mean pixel intensity was performed on IMARIS imaging software by generating a pixel mask for the green fluorescence channel after setting a suitable threshold to negate background fluorescence.

#### Radiolabeled ^3^H-glutamine uptake

Briefly, L-[3,4^3^H(N)]-glutamine (^3^H-glutamine; PerkinElmer, #NET551001MC) uptake was carried out using 1 million cells resuspended in 0.4 mL HBSS (Thermo, #14025092) containing ^3^H-glutamine (0.5 μCi/mL) and layered over 0.5 mL of 1:1 silicone oil:dibutyl phthalate (Sigma-Aldrich, #175633 and #524980 respectively). Uptake time was 3 mins, after which cells were pelleted below the oil via centrifugation, stopping uptake. The aqueous supernatant solution, followed by the silicone oil/dibutyl phthalate mixture, was aspirated and the cell pellet resuspended in 200 μL of 1 M NaOH. β-radioactivity was measured by liquid scintillation counting in a Beckman LS 6500 Multi-Purpose Scintillation Counter (scintillant Optiphase HiSafe 3; PerkinElmer, #1200.437). L-Alanine (Sigma, #A7627), L-Glutamine (Sigma, #G8540), 2-Amino-2-norbornanecarboxylic acid (BCH; Sigma, #A7902), α-(Methylamino)isobutyric acid (MeAIB; Sigma, #M2383), AHA or HPG were used at titrating concentrations or 5 mM to inhibit radiolabelled ligand uptake as indicated in the graphs.

## Notes

### Competing Interest Statement

The authors have declared no competing interest.

